# Benchmarking miniaturized microscopy against two-photon calcium imaging using single-cell orientation tuning in mouse visual cortex

**DOI:** 10.1101/494641

**Authors:** Annet Glas, Mark Hübener, Tobias Bonhoeffer, Pieter M. Goltstein

## Abstract

Miniaturized microscopes are lightweight imaging devices that allow optical recordings from neurons in freely moving animals over the course of weeks. Despite their ubiquitous use, individual neuronal responses measured with these microscopes have not been directly compared to those obtained with established *in vivo* imaging techniques such as bench-top two-photon microscopes. To achieve this, we performed calcium imaging in mouse primary visual cortex while presenting animals with drifting gratings. We identified the same neurons in image stacks acquired with both microscopy methods and quantified orientation tuning of individual neurons. The response amplitude and signal-to-noise ratio of calcium transients recorded upon visual stimulation were highly correlated between both microscopy methods, although influenced by neuropil contamination in miniaturized microscopy. Tuning properties, calculated for individual orientation tuned neurons, were strongly correlated between imaging techniques. Thus, neuronal tuning features measured with a miniaturized microscope are quantitatively similar to those obtained with a two-photon microscope.

## Introduction

In recent years, the arsenal of imaging techniques for neuroscience has been supplemented with miniaturized microscopes, of which several versions are currently available [1–3]. Miniaturized microscopes allow simultaneous, functional imaging of hundreds of neurons in a variety of brain areas in freely moving animals as small as a mouse over extended periods of time [2, 4, 6]. Key merits of miniaturized microscopes as compared to benchtop microscopes are the ability for head-mounting and their low cost [1]. These qualities make miniaturized fluorescence microscopy a valuable complementary method to other *in vivo* imaging techniques [2]. A trade-off compared to two-photon microscopes is the lack of optical sectioning, resulting in poorer lateral and axial resolution due to out-of-focus fluorescence. In addition, conventional miniaturized microscopes have a reduced ability for imaging deeper in the tissue, which is inherent to single-photon versus two-photon illumination wavelengths [5]. Together, these factors prevent *in vivo* imaging of sub-cellular structures such as dendritic spines as of yet [2]. On the positive side, miniaturized microscopy does enable chronic imaging of neurons and circuits in behavioral paradigms that require minimally constrained movement of the animal, and it has even been used as an alternative to functional two-photon imaging in head-fixed paradigms [7].

Despite the increasing use of miniaturized microscopy, signal amplitudes and neuronal tuning properties obtained with miniaturized microscope imaging have not been directly compared to those assessed with established *in vivo* imaging methods. Receptive field properties of neurons in primary visual cortex (V1) provide a suitable model for a direct comparison between both methods. Responses of visual cortex neurons to drifting gratings of particular orientations have been extensively investigated (e.g. [8, 9]). Individual neurons respond selectively to gratings of particular orientations and their preferred orientation remains largely stable across longer periods of time [9–12]. Here, we perform *in vivo* miniaturized and two-photon microscopy of neurons in V1 of anesthetized mice presented with moving gratings. We identify the same neurons with both microscopy techniques, and quantify the similarity in response properties of matched neurons.

## Materials and methods

### Animals

All procedures were performed in accordance with the institutional guidelines of the Max Planck Society and the local government (Regierung von Oberbayern, Germany). Eight female C57BL/6J mice (∼P60 on day of surgery) were individually housed in ventilated cages and kept on an inverted 12-h light, 12-h dark cycle with lights on at 10 AM. Ambient temperature (∼22°C) and humidity (∼55%) were kept constant. Water and standard chow were available ad libitum.

### Surgery

Mice were anesthetized with a mixture of fentanyl, midazolam, and medetomidine (FMM; 0.05 mg kg^−1^, 5 mg kg^−1^, and 0.5 mg kg^−1^ respectively, injected i.p.) and depth of anesthesia was monitored throughout the procedure by observation of the breathing rate and absence of a pedal reflex. Mice were placed in a stereotaxic apparatus (Neurostar) equipped with a thermal blanket (Harvard Apparatus). Eyes were covered with a thin layer of ophthalmic ointment. Lidocaine (0.2 mg ml^−1^ was sprayed onto the scalp for topical analgesia and carprofen (5 mg kg^−1^, injected s.c.) was administered for analgesia. The skull was exposed, dried and scraped with a scalpel. A custom-designed aluminum head bar was positioned using cyanoacrylate glue and subsequently covered with dental acrylic (Paladur). The location of V1 was verified using intrinsic signal imaging [22, 23] and a 4 mm circular craniotomy was created centered over V1. To sparsely label a population of V1 excitatory neurons, mice were injected with a viral vector mixture consisting of AAV2/1 CamKII0.4-Cre (1.15·10^10^ GC ml^−1^, Penn Vector Core) and AAV2/1 hSyn-flex-GCaMP6s (7.26·10^12^ GC ml^−1^, Penn Vector Core). At each injection site, 125 nl of viral vector was injected using a beveled glass pipette (30 *µ*m outer diameter) at an injection speed of 25 nl min^−1^. The glass pipette was slowly retracted 10 min after initial placement. Upon injection, the craniotomy was covered with a circular cover glass (4 mm, Warner Instruments), which was glued in place using cyanoacrylate gel and subsequently cemented with dental acrylic. After surgery, mice were injected with a mixture of antagonists (naloxone, flumazenil, and atipamezole; 1.2 mg kg^−1^, 0.5 mg kg^−1^, and 2.5 mg kg^−1^ respectively, injected s.c.) and left to recover under a heat lamp. Carprofen (5 mg kg^−1^, injected s.c.) was given on the three following days. Imaging experiments were conducted at least two weeks after surgery.

### Visual stimulation

Visual stimuli were displayed on a single LCD monitor (Dell P2717H; resolution: 1920 × 1080 pixels, width 60 cm, height 34 cm), with the center placed at roughly 45° azimuth and 12 cm from the animal’s eye. To assess orientation tuning, we presented full-screen square wave gratings (8 directions, 45° spacing) with a spatial frequency of 0.04 cycles per degree and a temporal frequency of 1.5 Hz. The stimulus set was flanked with a 30 s pre- and post-stimulation period. Each trial consisted of 3 s of moving grating, followed by 5 s of inter-trial interval during which a gray screen was presented. During both miniaturized and two-photon microscopy imaging sessions, the complete stimulus set was repeated five times, with a random order of directions in each repetition (trial). To avoid stimulus light leak during two-photon imaging, monitor illumination was shuttered during each scan-line and only turned on during the line-scanner turnaround period [24]. The space between the microscopy objective and cranial window was closed off using opaque tape.

### Miniaturized microscopy

Images were acquired with a commercially available miniaturized microscope (Basic Fluorescence Microscopy System - Surface, Doric Lenses) at a frame rate of 20 Hz and a resolution of 630 × 630 pixels (field of view 1 × 1 mm). Laser power under the objective lens (2× magnification, 0.5 NA) was <1 mW for all imaging experiments. The excitation wavelength was 458 nm. To minimize movement, the miniaturized microscope was mounted on a rigid holder (Doric Lenses) attached to an XYZ translation stage (Luigs Neumann).

For miniaturized microscope imaging experiments, mice were anesthetized with FMM (0.04 mg kg^−1^, 4 mg kg^−1^, and 0.4 mg kg^−1^ respectively, injected i.p.) The miniaturized microscope was positioned above the cranial window and lowered until the cortical surface blood vessel pattern became visible. To facilitate identification of individual neurons across microscopy techniques, a 3-dimensional volume spanning a depth 250 *µ*m was acquired at 5 *µ*m intervals between imaging planes while no visual stimulus was presented. Subsequently, visual stimuli (see above) were presented during imaging. Per session, up to 10 imaging planes were recorded in layer 2/3 at 10 *µ*m depth intervals. The onset of imaging was approximately 60 minutes after the administration of anesthesia, and the total duration of recording was typically under 75 minutes.

### Two-photon microscopy

Two-photon imaging was performed on a custom-built two-photon laser-scanning microscope with a Mai Tai eHP Ti:Sapphire laser (Spectra-Physics) set to a wavelength of 910 nm and a Nikon water immersion objective (16× magnification, 0.8 NA). Images were acquired with an image resolution of 750 × 800 pixels at a frame rate of 10 Hz. The field of view for functional imaging was 300 × 320 *µ*m. Laser power under the objective was kept stable at 25 mW throughout the experiment. Imaging data were acquired using custom software written in LabVIEW (National Instruments).

Two-photon imaging experiments were conducted one week after the miniaturized microscope imaging session in one half of the animals (n = 4 mice) and one week before the miniaturized microscope imaging session in the other half (n = 4 mice). Mice were anesthetized with FMM (0.04 mg kg^−1^, 4 mg kg^−1^ and 0.4 mg kg^−1^ respectively, injected i.p.). The imaging location of the previous miniaturized microscopy session was determined by comparing the blood-vessel pattern using a wide-field camera that was aligned with the two-photon microscope (Teledyne DALSA Inc.). Subsequently, the matched field of view was imaged with the two-photon microscope. Prior to functional imaging, a volume of 300 × 320 × 200 *µ*m (XYZ) was imaged at 1 *µ*m intervals while no visual stimulus was presented. For functional imaging, the anesthetized animal was presented with visual stimuli (see above), repeated for up to six imaging planes with depth increments of 10 *µ*m.

### Immunohistochemistry

Mice were deeply anesthetized and transcardially perfused with 9.25% w/v sucrose in distilled water followed by 4% PFA in PBS. Brains were then dissected out and post-fixed in 4% PFA for one week at 4°C. Coronal sections (50 *µ*m) were cut on a microtome (Thermo Fisher Scientific) and were kept free-floating at 4°C until further processing. Immunohistochemistry was carried out using the primary antibodies chicken anti-GFP (1:1000; Millipore) labeling GCaMP6s and rabbit anti-Homer3 (1:250; Synaptic Systems), which labels excitatory neurons. After washing, sections were incubated with species-specific secondary antibodies conjugated to Alexa Fluor 488 (1:200; Life Technologies) or Cy3 (1:200; Life Technologies) and mounted with mounting medium containing DAPI (Vectashield).

Images were acquired using a laser-scanning confocal microscope (Leica, TCS SP8), across serial optical sections (spaced at 1 *µ*m) acquired with a 20× objective (NA 0.75) at a resolution of 1024 × 1024 with a sequential scan using excitation lasers for DAPI (405 nm), Alexa488 (488 nm) and Cy3 (561 nm). Quantitative analysis was performed with the “Cell Counter” plug-in for ImageJ, by counting GFP expressing cells among Homer 3 expressing cells in cortical layer 2/3 (100-300 *µ*m from the pial surface).

### Analysis

Analysis of imaging data was performed using custom written routines in Matlab R2016b (Mathworks) and manual routines in the Fiji package of ImageJ (US National Institutes of Health) [25]. Small in-plane movement artefacts were corrected by aligning the images to a template [26]. Next, to identify the same neurons imaged with both microscopes, images obtained with both microscopes were scaled to match pixel size and image orientation. Initial alignments were made based on the cell location relative to major landmarks such as blood vessels. Once two pairs of neurons were judged to be identical in both imaging planes, the images were aligned using an ImageJ plugin (Align Image by line ROI). Subsequently, other cell pairs were identified based on absolute distance relative to other cells and blood vessels.

Cellular fluorescence signals were calculated for each imaging frame by averaging across all pixels within manually drawn regions of interest (ROIs). Fluorescence signals from miniaturized microscope recordings were corrected for local neuropil contamination by subtracting the average fluorescence in a 27 *µ*m ring (using a neuropil correction factor of 1.0) [13]. Because of the sparse labelling with GCaMP6, neuropil subtraction was not necessary for data acquired with the two-photon microscope (see also Results and Fig. 2e). In addition, for a small subset of data constrained non-negative matrix factorization for endoscopic data (CNMF-E) [14] was applied to miniaturized microscopy imaging frames, using a Gaussian kernel width 3.17 *µ*m, maximum soma diameter 23.8 *µ*m, minimum local correlation 0.8, minimum peak-to-noise ratio 6, spatial overlap ratio 0.05 and temporal correlation 0.8. For comparison of calcium transients, we determined putative sources to be identical between the methods when the CNMF-E-detected seed pixel and manually detected center pixel were less than 10 *µ*m apart, and if we could visually confirm similarity of the detected ROI contours. Next, ΔF/F calcium signals were quantified as relative increase in fluorescence over baseline, which was derived from the mean lowest 50% values in a 60 s sliding window [27]. In order to compare signal and noise amplitudes, miniaturized microscope data were resampled to the frame rate of the two-photon microscope (10 Hz). For each neuron, the signal amplitude was determined as the largest mean (across trials) response to any of the eight visual stimuli. Noise amplitude was calculated as the standard deviation of the ΔF/F values in the two-second period before stimulus presentation.

A neuron was defined as orientation tuned if it matched two criteria. First, the response to any of the eight movement directions was significantly different from any of the other directions (p<0.01), tested using the non-parametric Kruskal-Wallis test. Second, we excluded neurons of which the response to the preferred direction did not exceed the median response amplitude of the entire population of neurons (miniature microscope: 0.115 ΔF/F; two-photon microscope: 0.0525 ΔF/F; see Results and Fig. 3b). Orientation tuning curves were constructed by averaging the response to each movement direction and fitted with a two peaked Gaussian curve [28]. Preferred orientation was defined as the maximum of the fitted curve, and the tuning curve bandwidth was defined as half width of the fitted curve at 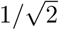 maximum. To quantify global orientation selectivity, we determined the normalized length of the mean response vector (also referred to as 1-circular variance or 1-CV) [29].

### Statistics

Normality of distributions was verified using the Kolmogorov-Smirnov test. Similarity of two different distributions was analyzed with the two-sample Kolmogorov-Smirnov test. A Mann-Whitney U test was used to compare distribution medians (Mdn). The tuning features circular variance and bandwidth of individual neurons were compared by computing the Spearman correlation coefficient r_s_ between both microscopy techniques and the preferred orientation was compared using the circular correlation coefficient r_circ_ (The orientation space was remapped to the range of 0 to 2*π* for this purpose) [30]. 95% confidence intervals of the median (Fig. 2e) were calculated using bootstrap resampling (bootstrap sample size: 84, number of re-samples: 10000). For all statistical tests, alpha was set at 0.05 and tests were conducted two-tailed.

## Results

### Identifying the same neurons across microscopy techniques

To compare evoked neuronal responses as obtained with a commercially available miniaturized microscope (Doric Lenses) and a custom-built two-photon microscope, we imaged V1 excitatory layer 2/3 neurons expressing the genetically encoded calcium indicator GCaMP6s while the anesthetized mice (n = 8) were presented with drifting square wave gratings (Fig. 1a) [11]. To minimize recording fluorescence from out-of-focus neurons, we used a dual viral vector intersectional approach and reduced the titer of the Cre-expressing viral vector, resulting in sparse labelling of neurons (see Methods; Fig. 1b). Post-hoc immunohistochemical analysis revealed that 32.6 ± 7.3 % of excitatory layer 2/3 neurons were labelled in the core of the bolus (injection titer of 1.15·10^10^ GC ml^−1^; Fig. 1c). Using superficial blood vessels as landmarks, we centered the fields of view of both microscopes on the same location. To overcome the differences in optical sectioning of both microscopes, we compared a single, background-subtracted field of view recorded with the miniaturized microscope with a two-photon microscope stack, collapsed along the axial axis spanning a depth of 100 *µ*m (Figs. 1d, e). Upon completion of both imaging sessions, neurons were matched based on their position relative to blood vessels, other identified neurons and relative depth in the tissue (Figs. 1d, e).

**Fig 1.**
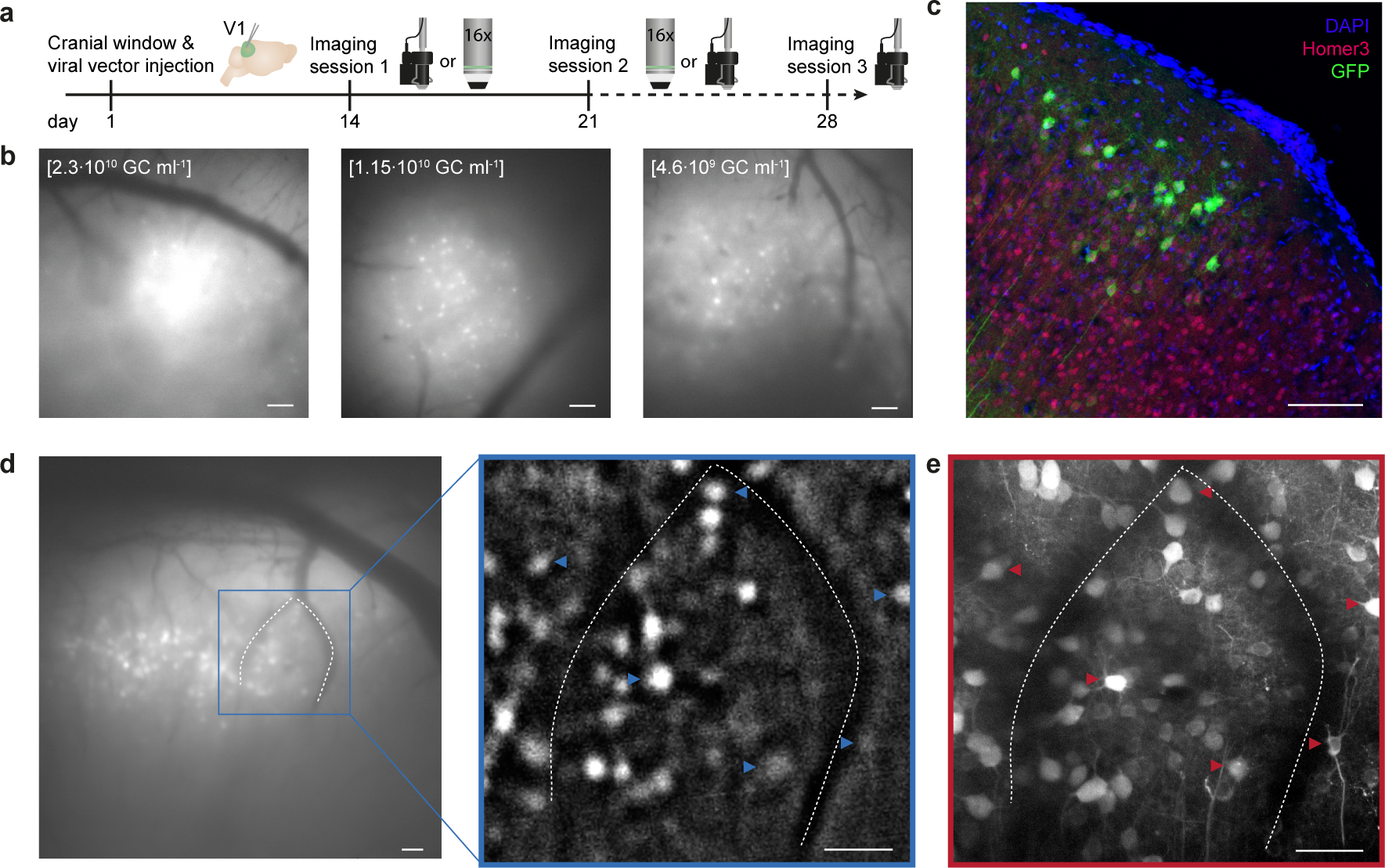
Matching neurons in images acquired with a miniaturized microscope and a two-photon microscope. (a) Experimental timeline in days. Day 1: Cranial window implant and viral vector injection into visual cortex layer 2/3. Days 14 and 21: One miniature and one two-photon microscopy imaging session on either day (order was counterbalanced across animals). Day 28: Optional second miniature microscopy session. Icons at day 14, 21 and 28 illustrate a miniature microscope and a 16× two-photon microscope objective. (b) Example miniaturized microscopy images of V1 injected with different viral vector titers (left: 2.3·10^10^ GC ml^−1^; middle: 1.15·10^10^ GC ml^−1^; right: 4.6·10^9^ GC ml^−1^). (c) Immunohistochemical labeling of GCaMP6s-expressing excitatory layer 2/3 neurons (injection titer of 1.15·10^10^ GC ml^−1^). (d) Left: Miniaturized microscopy image prior to processing. Right: Magnified image after background-subtraction. Blood vessels (dotted lines) assist in matching neurons between microscopes (see panel e; examples of matched neurons are indicated with arrowheads). (e) A collapsed volume as imaged with the two-photon microscope (100 planes, 1 *µ*m spacing, projection along the axial axis). Scale bars, 100 *µ*m (b, c) and 50 *µ*m (d, e).

### Extraction of stimulus-evoked responses of matched neurons

After careful, off-line matching of neurons across images, we were able to identify 488 neurons that were present in both fields of view (Fig. 2a). This matching method allowed us to recognize a match for many, but not all neurons in the imaged planes. Of note, we found that the population of neurons detected in a single miniaturized microscopy imaging plane spanned over 70 *µ*m in depth within the two-photon imaged volume (Fig. 2b).

**Fig 2.**
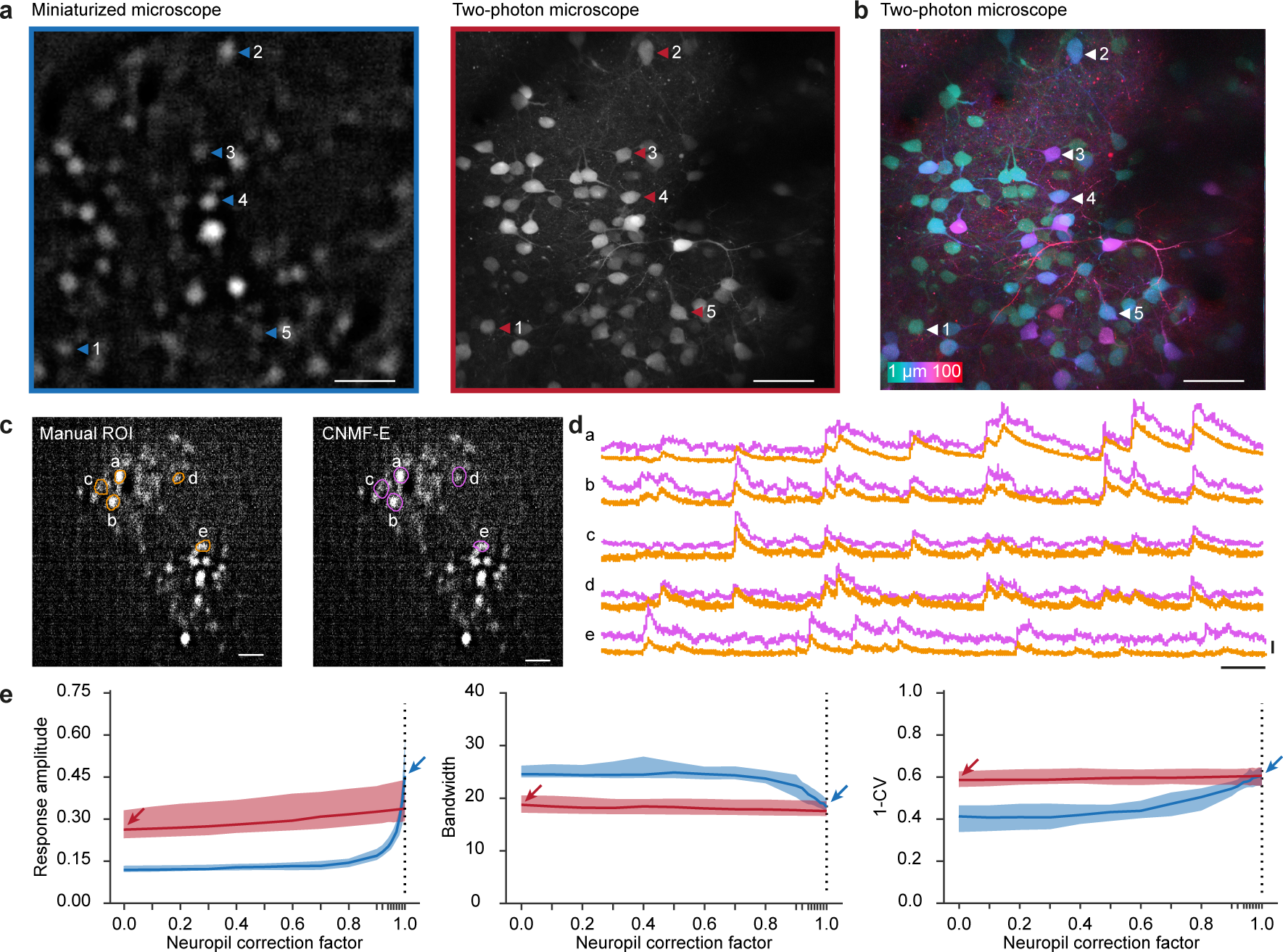
Imaging of visually evoked responses using a miniaturized microscope and a two-photon microscope. (a) A background-subtracted image acquired with a miniaturized microscope (left) and a collapsed volume (100 planes, 1 *µ*m spacing) acquired with a two-photon microscope (right). Example neurons matched across microscopes are indicated with arrowheads. (b) Pixel-wise color-coded depth origin of the collapsed two-photon volume. (c) Contours of neurons detected with either manual ROI selection (orange, left) or CNMF-E (pink, right) within the same background-subtracted field of view recorded with a miniaturized microscope. (d) Relative fluorescence changes (ΔF/F) of example neurons indicated in (c). (e) Median response amplitude (ΔF/F), bandwidth and global orientation selectivity index (1-CV) as a function of neuropil correction factor in recordings acquired from orientation tuned neurons using a miniaturized microscope (blue) and a two-photon microscope (red). Arrows indicate parameter values at the selected neuropil correction factor of miniature microscopy (blue) and two-photon microscopy (red). Colored shading indicates 95% confidence interval. Scale bars, 50 *µ*m (a, b), 100 *µ*m (c), 25 s (d, horizontal), 1 ΔF/F (d, vertical).

Before analyzing calcium activity from these neurons, we explored whether the obtained calcium transients were robust to the choice of signal extraction method. To this end, we extracted single cell calcium transients in a subset of miniaturized microscopy recordings by manual region of interest (ROI) selection [13], which involves outlining of the neurons’ contours and direct surrounding by the experimenter (see Methods). We contrasted this to constrained nonnegative matrix factorization for microendoscopic data (CNMF-E) [14, 15], which decomposes the recorded fluorescence into spatial footprints and temporal components modelling the calcium dynamics (Figs. 2c,d). The obtained calcium transients were similar in both signal amplitude and transient kinetics (Fig. 2d). However, source extraction methods often return spatial footprints that extend beyond the boundaries of visually identified neurons and tend to ignore cells that show only little calcium activity. Because re-identification of neurons across microscopy techniques relied on a direct comparison of morphological information, independent of calcium activity, we chose to quantify calcium traces and tuning properties using manual ROI selection.

An important step in the calculation of single cell calcium traces using manual ROI selection is to correct the contamination of cellular signals by out-of-focus fluorescence from the neuropil. The method, referred to as neuropil correction [13], measures neuropil fluorescence from an area directly surrounding the cell and subtracts the neuropil-signal time course, scaled by a factor (neuropil correction factor), from the signal measured within the outline of the cell. The rationale is that the signal measured within the cellular ROI is the linear sum of two signals: one truly generated in the ROI and one originating from tissue adjacent to the ROI (due to the limited axial and/or lateral resolution as well as tissue scattering). By choosing an appropriate neuropil correction factor, the contamination can be corrected by subtraction of the scaled neuropil time course. In our experiments, labeling of cortical cells was sparse, and we observed only a very small amount of neuropil fluorescence in the two-photon microscopy recordings (Fig. 2a, right panel). Hence, we chose to use a neuropil correction factor of 0.0 for these experiments. In contrast, miniaturized epifluorescence microscopy lacks optical sectioning, therefore we assumed that in those experiments virtually all of the signal originating from above and below an outlined neuron will mix into the measured neuronal signal, resulting in an estimated neuropil correction factor of 1.0.

In order to test whether our choice of neuropil correction factor for each method was appropriate, we varied the neuropil correction factor from 0.0 to 1.0 and investigated how three key parameters in this study changed as a result (the parameters were response amplitude, bandwidth and the global orientation selectivity index 1-circular variance (1-CV) of orientation tuned cells; see below and Methods for further explanation). The analysis showed that, in our two-photon microscopy recordings, these parameters altogether depended very little on the choice of neuropil correction factor (Fig. 2e). This indicated that neuropil contamination was negligible, and it validated the choice for the value of 0.0 in two-photon microscopy recordings. However, in miniaturized microscopy recordings, all three parameters depended strongly on the neuropil contamination factor; signal amplitude and orientation selectivity (1-CV) increasing monotonically and bandwidth decreasing monotonically (Fig. 2e). The curves for miniaturized and two-photon microscopy recordings intersected when the neuropil correction factor approximated the maximum value of 1.0, suggesting that the choice of neuropil correction factor in miniaturized microscopy recordings (1.0) is close to the optimal value.

### Orientation tuned neurons show similar tuning properties in miniaturized microscope and two-photon microscope recordings

We extracted the calcium transients of all matched neurons and quantified the responses to visual stimulation (Fig. 3a). Both the average response amplitude (r_s_ = 0.602, p = 1.841·10^−49^, n = 488 neurons; Fig. 3b) and the ΔF/F signal-to-noise ratio (r_s_ = 0.407, p = 6.348·10^−21^, n = 488 neurons; Fig. 3b) measured using the miniaturized microscope correlated strongly with the measurements recorded using the two-photon microscope. The median visually evoked response amplitude was significantly higher in miniaturized microscopy recordings (Mdn = 0.115) compared to two-photon microscopy recordings (Mdn = 0.0525, Wilcoxon test, T = 92271, p = 1.268·10^−25^).

**Fig 3.**
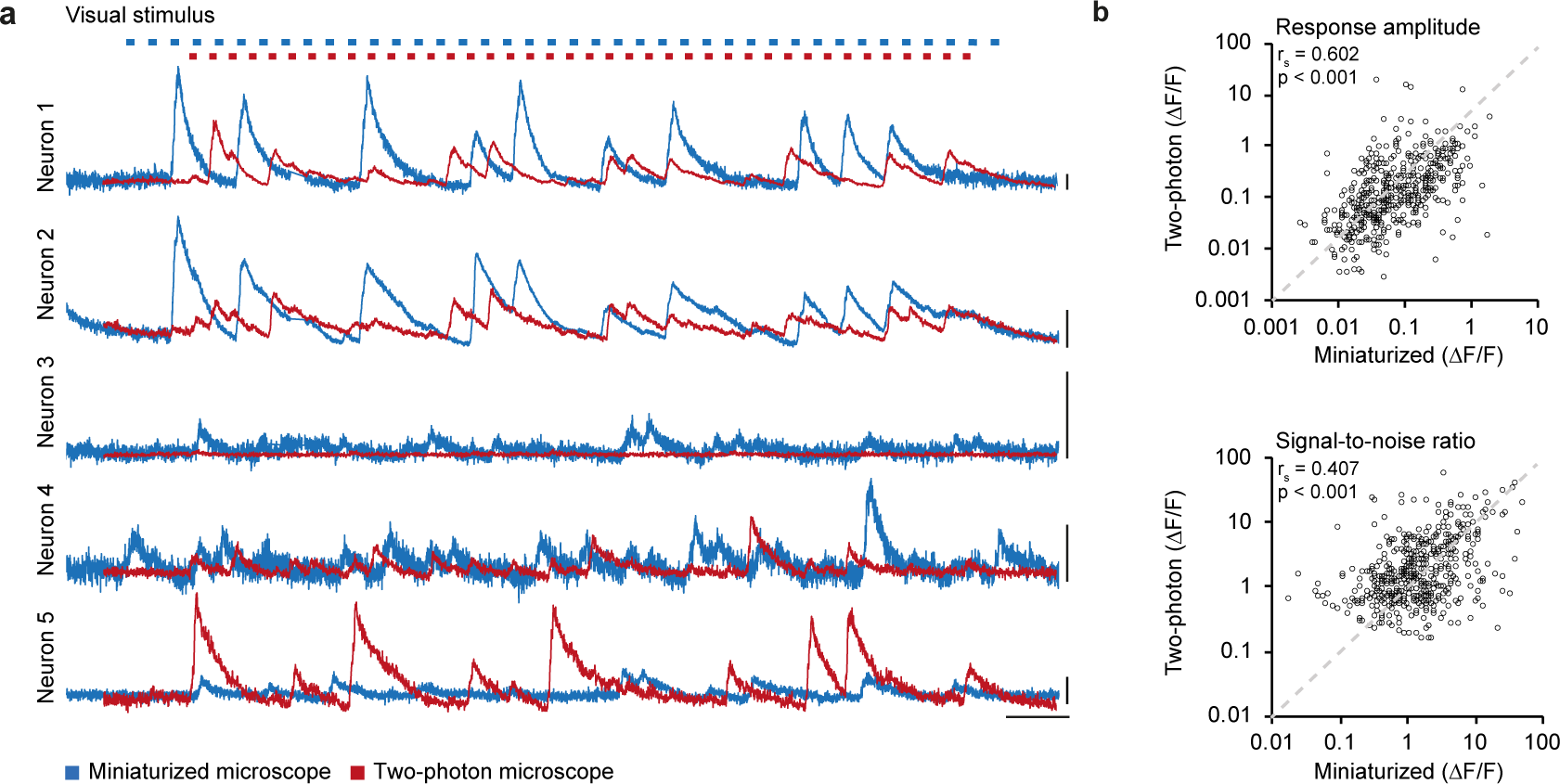
Calcium traces in miniaturized microscope and two-photon microscope recordings. (a) Relative fluorescence changes (ΔF/F) for the five example neurons depicted in Fig. 2a,b as recorded with a miniaturized microscope (blue) and a two-photon microscope (red) during visual stimulation. Top: Blue and red marks indicate stimulus presentation for miniaturized microscopy and two-photon microscopy, respectively. Stimuli were presented in a pseudo-randomized order that was unique for each experiment. (b) Average ΔF/F response amplitude to the preferred stimulus (p = 1.841·10^−49^, Spearman’s correlation) and ΔF/F signal-to-noise ratio of stimulus-induced calcium transients (p = 6.348·10^−21^, Spearman’s correlation) of all matched neurons (488 neurons, n = 8 mice) in miniaturized microscope recordings plotted against the respective values in two-photon microscope recordings. Scale bars, 1 ΔF/F (a, vertical), and 25 s (a, horizontal).

In contrast, the median ΔF/F signal-to-noise ratio was significantly higher in two-photon microscopy recordings (Mdn = 1.432) compared to miniaturized microscopy recordings (Mdn = 1.339, Wilcoxon test, T = 69005, p = 0.003). Thus, while single-neuron visually driven fluorescence changes were strongly correlated between microscopes, the absolute values of response amplitude and signal-to-noise ratio were slightly different (this difference varies as function of the value of the neuropil correction factor; see Discussion and Fig. 2e).

Many V1 neurons respond preferentially to moving gratings of specific orientations (Fig. 4a) and their tuning features are relatively stable over the course of weeks [10, 12], making this response property ideally suited for a direct comparison of the microscopy techniques. We averaged the calcium responses to the moving gratings of different directions and fitted the responses with a two-peaked Gaussian curve (Fig. 4b). Out of 488 matched neurons, 194 were classified to be orientation tuned (see Methods) in miniaturized microscopy recordings, 133 in two-photon microscopy recordings, and 84 of these matched the criteria for being orientation tuned in both miniaturized and two-photon microscopy recordings (n = 7 mice; Fig. 5a).

**Fig 4.**
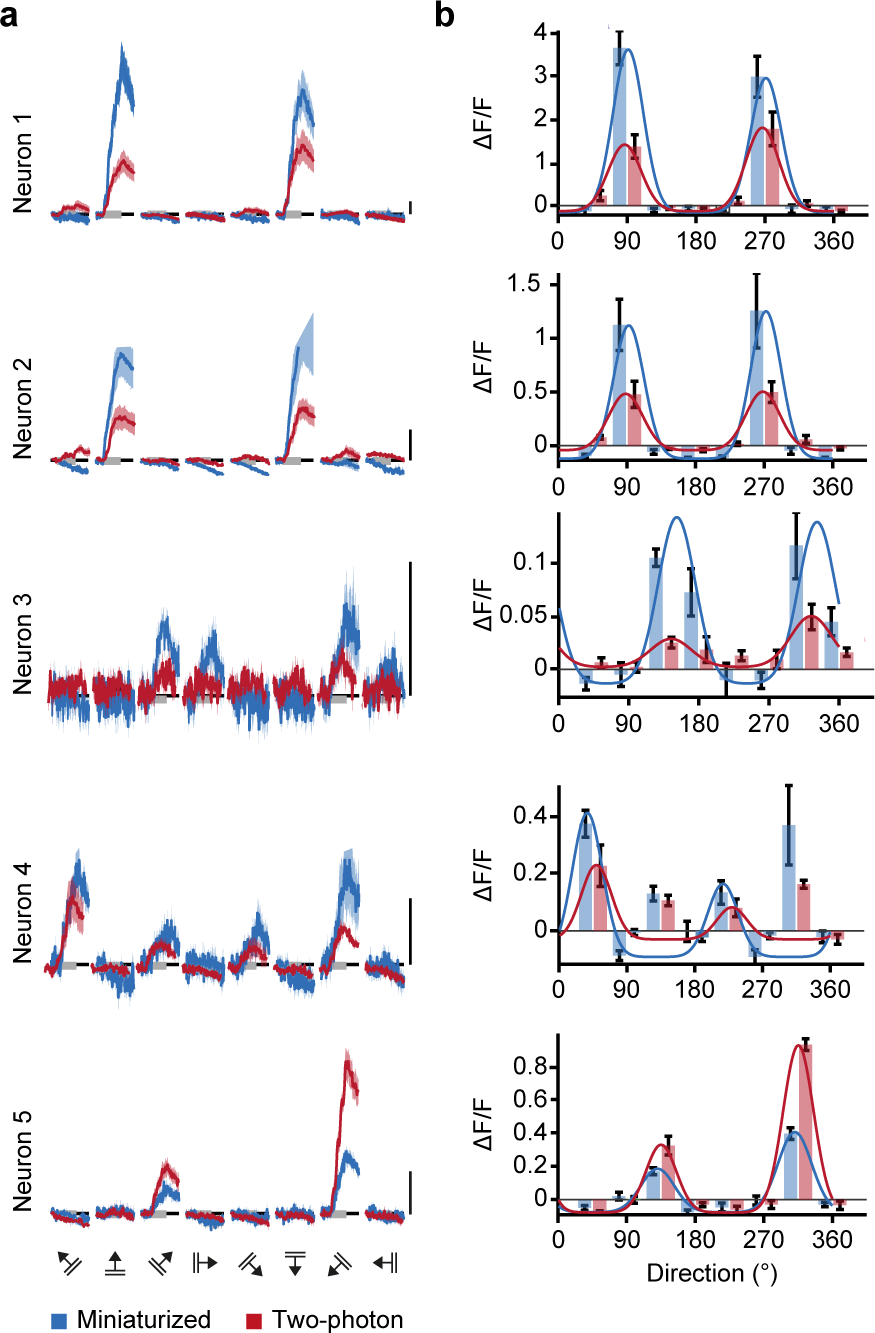
Orientation tuning curves of V1 neurons match between miniaturized microscopy and two-photon microscopy. (a) Calcium responses as acquired with a miniaturized microscope (blue) and a two-photon microscope (red) in response to drifting gratings (8 directions, 5 repetitions) of the five example neurons depicted in Fig. 2a,b. (b) Bars show the mean (± SEM) responses of the same neurons imaged with a miniaturized microscope (blue) and a two-photon microscope (red). Overlaid blue and red lines indicate the fitted tuning curves. Scale bars, 1 ΔF/F.

**Fig 5.**
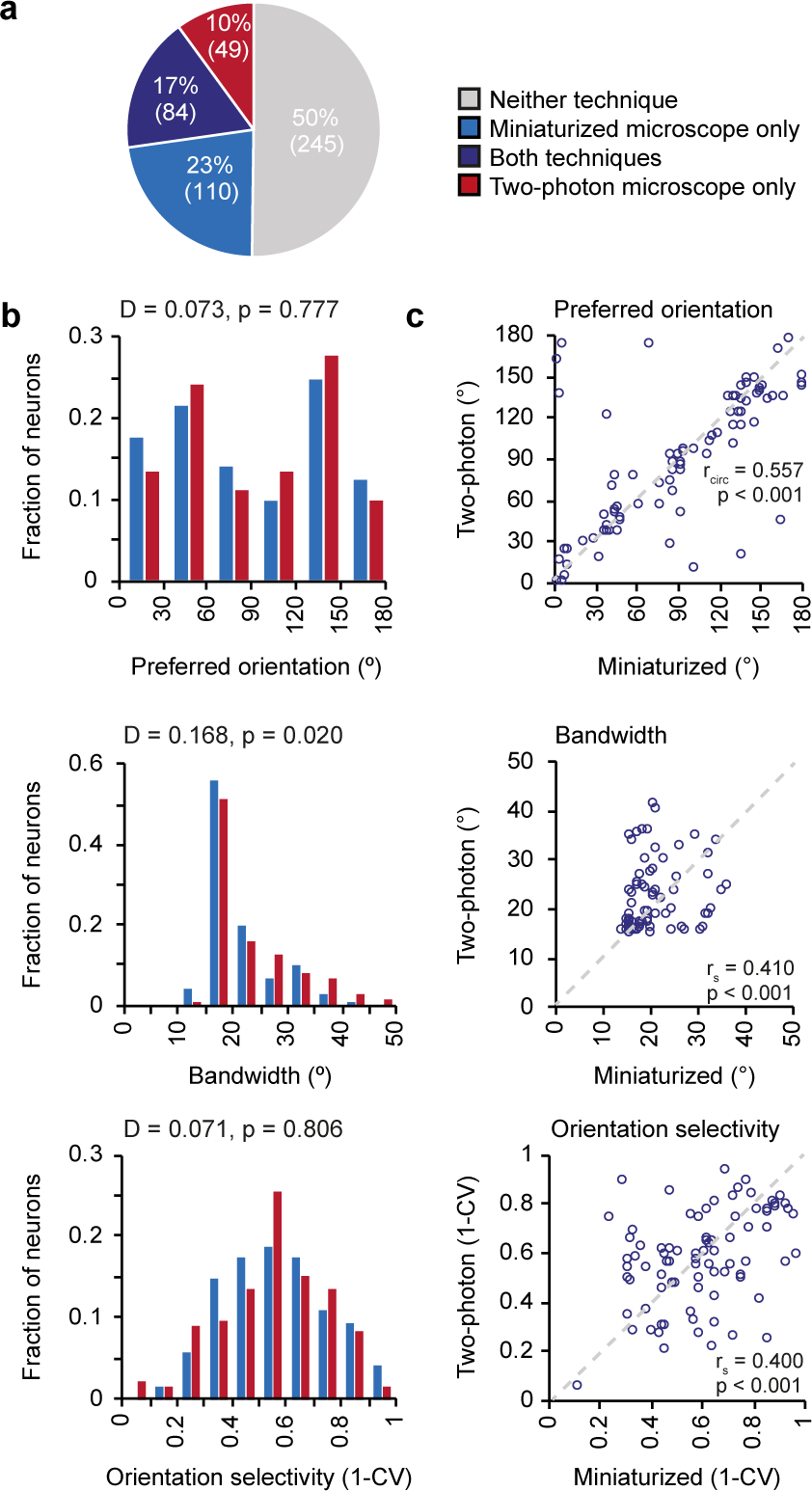
Orientation tuning properties of V1 neurons as imaged with a miniaturized microscope and a two-photon microscope. (a) Fractions of neurons that were classified as orientation tuned in miniaturized microscope recordings only (blue), using both microscopy techniques (purple), in two-photon microscope recordings only (red), or using neither microscopy technique (gray). (b) Distribution of preferred orientation (p = 0.777, two-sample Kolmogorov-Smirnov test), bandwidth (p = 0.020, two-sample Kolmogorov-Smirnov test), and global orientation selectivity index 1-CV (p = 0.806, two-sample Kolmogorov-Smirnov test) for all neurons that were orientation tuned in recordings with a miniaturized microscope (blue bars, 194 neurons, n = 7 mice) or a two-photon microscope (red bars, 133 neurons, n = 6 mice). (c) Preferred orientation (p = 2.02·10^−6^, circular correlation), bandwidth (p = 1.23·10^−4^, Spearman’s correlation), and global orientation selectivity index 1-CV (p = 1.81·10^−4^, Spearman’s correlation) for individual neurons (purple circles) that were orientation tuned using both microscopy techniques (84 neurons, n = 5 mice). The unity line is depicted as gray dashed line.

Key parameters of the tuning curves (preferred orientation, bandwidth, and global orientation selectivity, as described by 1-CV) were first determined for all neurons that were orientation tuned in each microscopy technique separately (Fig. 5b). The overall distributions for preferred orientation (two-sample Kolmogorov-Smirnov test, D = 0.073, p = 0.777) and 1-CV (two-sample Kolmogorov-Smirnov test, D = 0.071, p = 0.806) did not significantly differ between microscopy techniques (n_miniaturized_ = 194, n_two-photon_ = 133 in 7 mice). However, tuning curve bandwidth was distributed differently between microscopy techniques (two-sample Kolmogorov-Smirnov test, D = 0.168, p = 0.020). This observation indicates that orientation tuning in two-photon recordings appeared slightly broader, as evidenced by a larger median bandwidth (Mdn_miniaturized_ = 18.9, Mdn_two-photon_ = 19.8, Mann-Whitney U-test, U = 29919, p = 0.024), while median global orientation selectivity did not significantly differ (1-CV, Mdn_miniaturized_ = 0.564, Mdn_two-photon_ = 0.561, Mann-Whitney U-test, U = 32239, p = 0.61). However, the existence of a difference between the distribution-median of tuning curve parameters depends on fine-tuning of the neuropil correction factor (see above, Discussion and Fig. 2e).

To compare the tuning properties at the single neuron level, we limited the analysis to neurons that were classified orientation tuned with both microscopy techniques (n = 84 in 5 mice). Most importantly in the present context, the preferred orientation (r_circ_ = 0.557, p = 2.02·10^−6^), bandwidth (r_s_ = 0.410, p = 1.23·10^−4^) and the global orientation selectivity index (1-CV; r_s_ = 0.400, p = 1.81·10^−4^) of these individual neurons correlated significantly between recordings performed with both microscopes (n = 84 neurons; Fig. 5c). As already quantified for the overall population, the average response amplitude across these orientation tuned neurons was again significantly higher in the miniaturized microscopy recordings (Mdn = 0.457) than in two-photon microscopy recordings (Mdn = 0.265, Wilcoxon test, T = 2913, p = 4.89·10^−7^ n = 84 neurons). However, in this specific subset of neurons the ΔF/F signal-to-noise ratio was significantly higher in the two-photon microscopy recordings (Mdn = 7.137) compared to the miniaturized microscopy recordings (Mdn = 4.838, Wilcoxon test, T = 1066, p = 0.001, n = 84 neurons).

### The effect of between-session variability on signal amplitude and tuning properties

To test to which extent observed differences between microscopy techniques could be explained by test-retest variance, we performed a second miniaturized microscopy session spaced one week apart from the first miniaturized microscopy session in four mice (Fig. 1a). In order to allow a direct comparison between this analysis and the results described above, we only considered neurons that were also observed in the accompanying two-photon microscopy session. As expected, both average response amplitude (r_s_ = 0.585, p = 4.78·10^−18^, n = 181 neurons in 4 mice; Fig. 6a) and the ΔF/F signal-to-noise ratio (r_s_ = 0.647, p = 6.45·10^−23^, n = 181 neurons in 4 mice; Fig. 6a) were strongly correlated between the two miniaturized microscopy sessions. When comparing the response amplitude correlation of two consecutive miniaturized microscopy sessions with the correlation between two sessions using the two different microscopes (Fig. 3b versus Fig. 6a), the correlation coefficient between these groups was not significantly different (Fisher’s r-to-z transformation, z = 0.3, p = 0.382).

**Fig 6.**
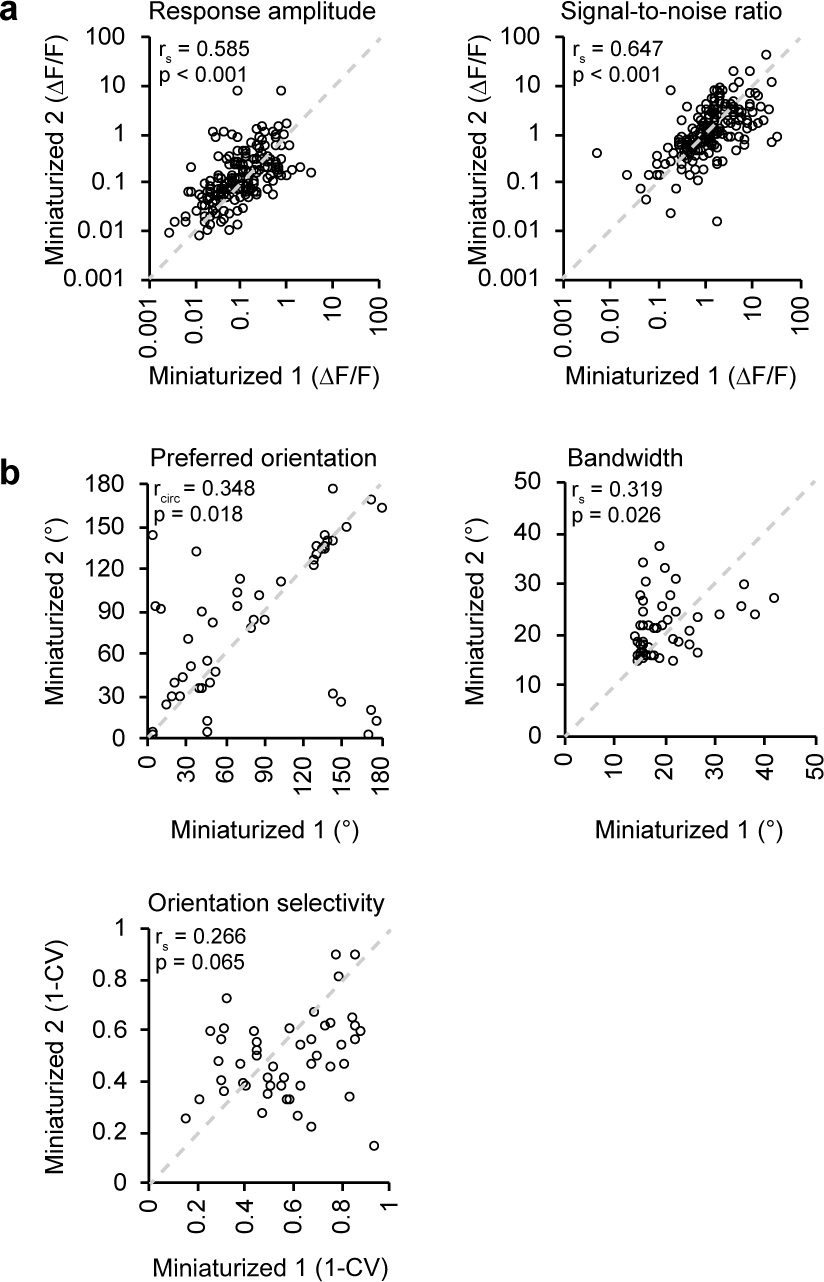
Effect of test-retest variability on recorded response properties in V1. (a) Average ΔF/F response amplitude to the preferred stimulus (p = 4.780·10^−18^, Spearman’s correlation) and ΔF/F signal-to-noise ratio of stimulation-induced calcium transients (p = 6.451·10^−23^, Spearman’s correlation) of matched neurons (181 neurons, n = 4 mice) in two consecutive miniaturized microscopy sessions (Miniaturized 1 and Miniaturized 2). (b) Preferred orientation (p = 0.018, circular correlation), bandwidth (p = 0.026, Spearman’s correlation), and global orientation selectivity index 1-CV (p = 0.065, Spearman’s correlation) for individual neurons (black circles) that were orientation tuned during both consecutive microscopy sessions and visually detected in the two-photon microscopy session (49 neurons, n = 3 mice).

Finally, we assessed tuning curve parameters of neurons that were orientation tuned in both miniaturized microscopy sessions, as well as visually detected in the two-photon microscopy session (n = 49 in 3 mice). The preferred orientation (r_circ_ = 0.348, p = 0.018) and bandwidth (r_s_ = 0.319, p = 0.026) of these individual neurons correlated significantly between test-retest conditions (Fig. 6b). However, the test-retest relationship between the global orientation selectivity index was not significant (1-CV; r s = 0.266, p = 0.065; Fig. 6b), possibly because of the low number of neurons that could be included in this analysis.

## Discussion

We used calcium imaging to measure visual response properties of V1 excitatory neurons with both a miniaturized microscope and a stationary two-photon microscope. The same neurons could be identified in images acquired with both microscopes. This was achieved by making use of sparse GCaMP6 labelling and volumetric structural imaging to overcome differences in optical sectioning between the two microscopy techniques. The amplitude and signal-to-noise ratio of visually evoked calcium transients of identical neurons were strongly correlated across imaging techniques and tuning features of orientation-tuned neurons recorded with the two microscopes were similar at the individual cell level. However, the population median of response and tuning parameters could be offset depending on the choice of neuropil correction factor that was applied to miniature microscopy data. The observed similarities were comparable to those between two consecutive miniaturized microscopy sessions. This suggests that the observed variability between microscopes is not larger than expected from miniature microscope test-retest variability. Overall, our results show that single-photon miniaturized microscopy is a reliable method for recording functional properties of neurons in the visual cortex.

### Influence of out-of-focus fluorescence

Although neuronal stimulus-induced calcium transients and orientation tuning features were strongly correlated between microscopy techniques at the single neuron level, we did observe certain differences when comparing the distributions of these features across the population of recorded neurons. The maximum amplitude of stimulus-induced calcium transients was larger in miniaturized microscope recordings, while the signal-to-noise ratio was lower. Furthermore, across the population of orientation-tuned neurons, local feature selectivity was slightly reduced as described by broader tuning curve bandwidths in recordings with the two-photon microscope.

Differences between the distributions of signal-to-noise ratio might be expected when comparing two imaging methods that differ vastly in the numbers of photons collected per neuron, e.g. using a CMOS sensor for miniaturized microscopy and a photomultiplier tube for two-photon microscopy. Moreover, differences in response amplitude and orientation selectivity can be attributed, at least in part, to the choice of the neuropil correction factor for analyzing miniaturized microscopy recordings. The curves describing the relationship between neuropil correction factor and ΔF/F response amplitude calculated for two-photon and miniaturized microscopy data intersect at a neuropil correction factor slightly smaller than 1.0 (see Fig. 2e). Empirically, it can therefore be argued that for miniaturized microscopy a neuropil correction factor slightly below 1.0 should be employed, which is also theoretically evident: the neuropil signal is estimated by calculating the mean of all fluorescence in the cell-devoid region directly adjacent to an ROI (e.g. a neuron). On the other hand, the measured neuronal signal is the sum of the true neuronal signal from the cell body and the neuropil signal originating from within the ROI, not including any neuropil signal from the axial/lateral range in which the neuron’s cell body was present. Thus, the intensity of neuropil signal bleeding into the neuronal signal is slightly lower than the intensity of neuropil signal measured in the area adjacent to the neuronal ROI. The optimal neuropil correction factor for an imaging method with poor optical sectioning (such as miniaturized microscopy) should therefore be just below 1.0, rather than exactly 1.0.

However, the empirically determined neuropil correction factor will depend on the density of neurons that expresses calcium indicator, and would have to be empirically verified for each preparation and tissue using a two-photon microscope, which is not practical for most studies. Therefore, for our purpose of verifying the general applicability of single cell calcium imaging using miniature microscopy, we think it is best to use the initial estimate of 1.0 as neuropil correction factor.

### Comparison of source extraction methods

A key feature of our approach is the direct matching of the same neurons between microscopy techniques. The two techniques differ considerably in their ability for optical sectioning, with an increased probability that two neurons, located at different depths, cannot be separated using manual annotation methods in miniaturized microscopy recordings. Therefore it was important to obtain a sparse population of labelled neurons, which we achieved by titrating down the Cre-expressing viral vector. To extract calcium signals from both miniaturized microscopy and two-photon microscopy data, we chose a conventional method for extracting ΔF/F calcium activity, which uses the mean fluorescence signal from manually detected ROIs. This method facilitated a direct, morphology-based comparison of individual neurons recorded with the two imaging techniques. However, there are alternative, activity-based automated ROI detection and source extraction methods that can be used for analyzing miniaturized microscopy and endoscopy data [14, 17]. These methods have the advantage of allowing to demix activity patterns of overlapping sources (cells) that are often observed in more densely labelled preparations. In a subset of miniaturized microscopy recordings, we show that the calcium transients detected by an alternative source extraction method, CNMF-E [14], are similar to those that we detected using our manual ROI approach. We therefore expect that our conclusions extend to the use of this (and similar) source extraction and deconvolution method(s) that allow for recordings with denser labelling than reported here.

### Session-to-session variability

Since the response properties of visual cortex cells are quite stable over time [10, 12, 18], we did not anticipate large differences in these properties to emerge within days. However, a portion of the variation in measured tuning properties between microscopy techniques might be ascribed to mere difference across time points, possibly relating to small fluctuations in anesthesia at the time of imaging. We conducted consecutive imaging sessions one week apart, with the first session performed two weeks after viral vector injection. We chose a one-week interval between imaging sessions to allow the animal to recover completely from anesthesia and to allow us to approximate the same anesthetic state in both experiments. Other forms of lightly dosed anesthesia, such as isoflurane, do not significantly alter V1 response properties as compared to awake animals [19]. However, we cannot exclude the possibility that fluctuations of fentanyl-based anesthesia can cause minor differences in orientation tuning between imaging sessions in our experiment. A study performed in awake experiments with minimal lag between imaging sessions might address these concerns but may at the same time suffer from other, e.g. state-dependent sources of inter-session variability [9, 20].

### Combining miniaturized and two-photon microscopy

The overall aim of our study was to quantitatively compare recordings obtained with miniaturized microscopy to those obtained with a conventional *in vivo* microscopy method such as two-photon microscopy. We report a high degree of similarity between these recordings, in spite of categorical differences between the two imaging methods [5]. A promising future approach would be to make use of both microscopy methods in a single experimental design, optimally using their respective qualitative merits. Such an approach could involve imaging of a population of neurons with a miniaturized microscope while an animal engages in a freely moving task and subsequently characterizing structural changes in neurons implicated in the task with a two-photon microscope. An exciting new possibility is two-photon miniaturized microscopy [21], which allows functional imaging of single dendrites and dendritic spines in freely behaving animals. However, the currently smaller field of view reduces the number of somata that can be imaged at once, which makes identification of (sparse) task-related neurons and of large-scale population activity dynamics challenging with this method. The combination of single-photon miniaturized microscopy and two-photon microscopy thus provides a promising approach to disentangle the processes at the functional and structural level that underlie behavior in freely moving animals.

## Acknowledgments

We thank Claudia Huber, Max Sperling, Volker Staiger, and Frank Voss for technical assistance.

## Author Contributions Statements

A.G. collected the data; A.G. and P.M.G. analyzed the data; all authors conceived the study and wrote the manuscript.

## Data Availability

The dataset of this study is available on

https://web.gin.g-node.org/pgoltstein/ Mini1p2pcomparison Glas Goltstein 2018.

